## Supplementary figures and images for "Rephine.r: a pipeline for correcting gene calls and clusters to improve phage pangenomes and phylogenies"

### Supplemental Fig. 1

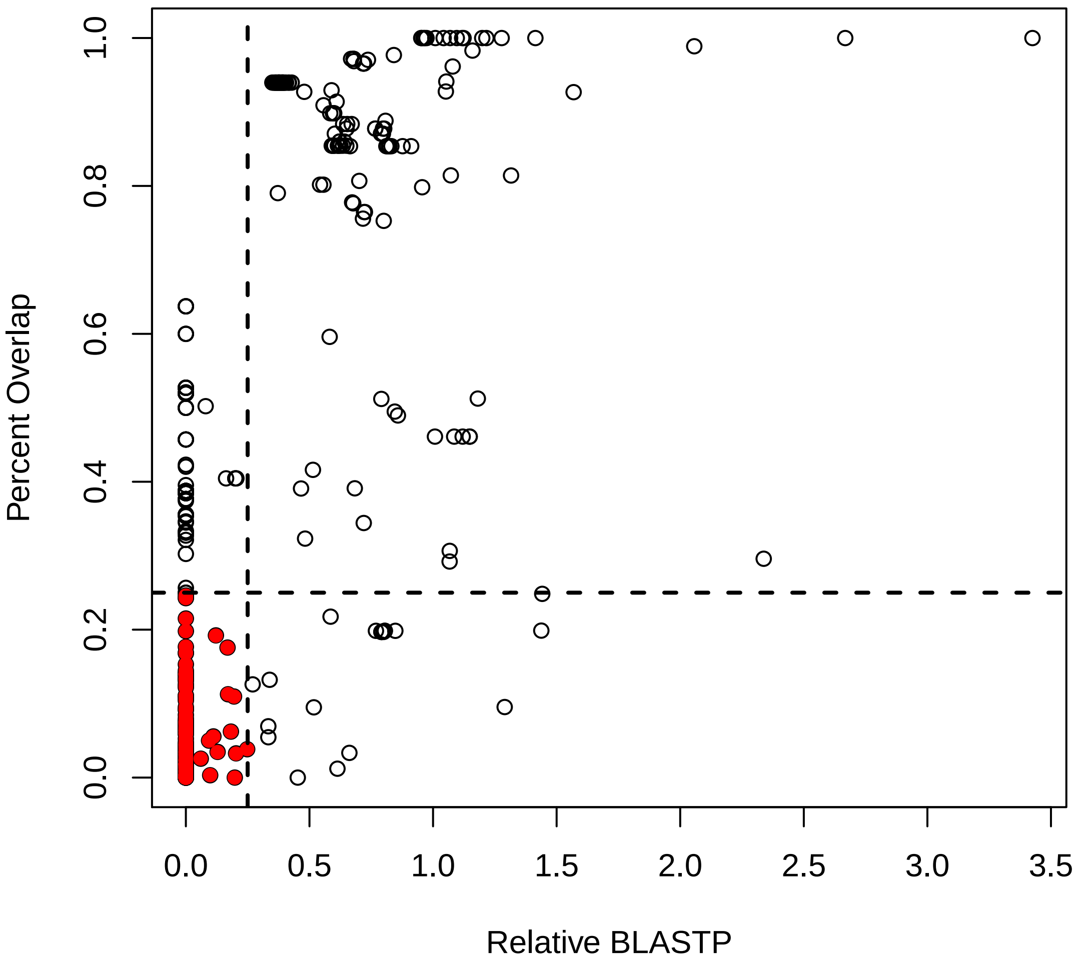

### Supplemental Fig. 2

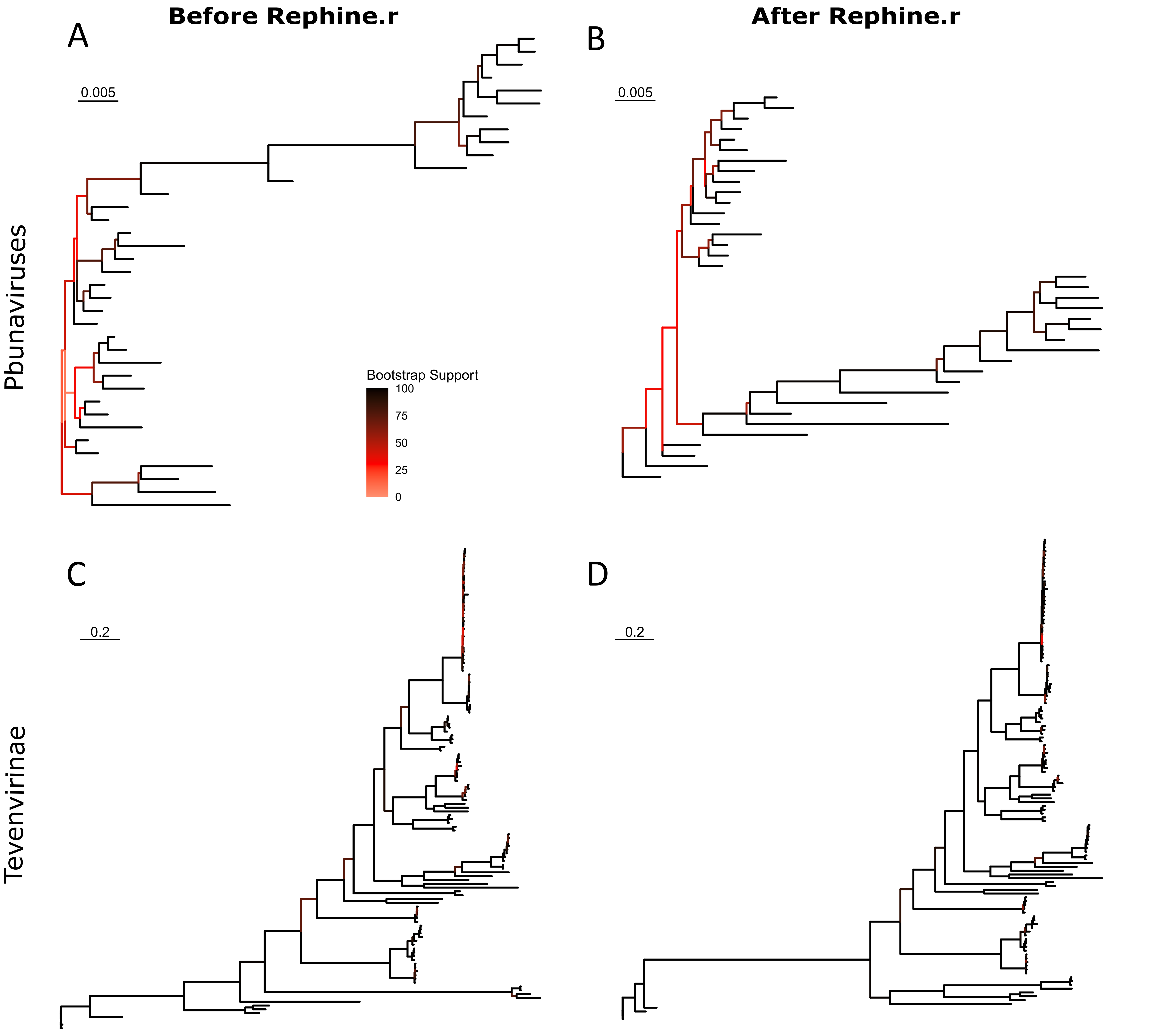
